# Dual metabolomic profiling uncovers *Toxoplasma* manipulation of the host metabolome and the discovery of a novel parasite metabolic capability

**DOI:** 10.1101/463075

**Authors:** William J. Olson, David Stevenson, Daniel Amador-Noguez, Laura J. Knoll

## Abstract

The obligate intracellular parasite *Toxoplasma gondii* is auxotrophic for several key metabolites and must scavenge these from the host. It is unclear how *Toxoplasma* manipulates host metabolism for its overall growth rate and non-essential metabolites. To address this question, we measured changes in the joint host-parasite metabolome over a time course of infection. Host and parasite transcriptomes were simultaneously generated to determine potential changes in metabolic enzyme levels. *Toxoplasma* infection increased activity in multiple metabolic pathways, including the tricarboxylic acid cycle, the pentose phosphate pathway, glycolysis, amino acid synthesis, and nucleotide metabolism. Our analysis indicated that changes in some pathways, such as the tricarboxylic acid cycle, derive from the parasite, while changes in others, like the pentose phosphate pathway, were host and parasite driven. Further experiments led to the discovery of a *Toxoplasma* enzyme, sedoheptulose bisphosphatase, which funnels carbon from glycolysis into ribose synthesis through a energetically driven dephosphorylation reaction. This second route for ribose synthesis resolves a conflict between the *Toxoplasma* tricarboxylic acid cycle and pentose phosphate pathway, which are both NADP+ dependent. During periods of high energetic and ribose need, the competition for NADP+ could result in lethal redox imbalances. Sedoheptulose bisphosphatase represents a novel step in *Toxoplasma* central carbon metabolism that allows *Toxoplasma* to satisfy its ribose demand without using NADP+. Sedoheptulose bisphosphatase is not present in humans, highlighting its potential as a drug target.

**Author Summary:** The obligate intracellular parasite *Toxoplasma* is commonly found among human populations worldwide and poses severe health risks to fetuses and individuals with AIDS. While some treatments are available they are limited in scope. A possible target for new therapies is *Toxoplasma*’s limited metabolism, which makes it heavily reliant in its host. In this study, we generated a joint host/parasite metabolome to better understand host manipulation by the parasite and to discover unique aspects of *Toxoplasma* metabolism that could serve as the next generation of drug targets. Metabolomic analysis of *Toxoplasma* during an infection time course found broad activation of host metabolism by the parasite in both energetic and biosynthetic pathways. We discovered a new *Toxoplasma* enzyme, sedoheptulose bisphosphatase, which redirects carbon from glycolysis into ribose synthesis. Humans lack sedoheptulose bisphosphatase, making it a potential drug target. The wholesale remodeling of host metabolism for optimal parasite growth is also of interest, although the mechanisms behind this host manipulation must be further studied before therapeutic targets can be identified.

## Introduction

*Toxoplasma gondii* is an obligate intracellular protozoan of the phylum Apicomplexa. It can infect any nucleated cell of any warm-blooded animal and forms a chronic infection in approximately one third of the human population [1]. Healthy human adults are often asymptomatic during infection, but *T. gondii* can be lethal to immunocompromised individuals and to fetuses if acquired congenitally [1]. As an obligate intracellular parasite *T. gondii* relies on host cell metabolism to compensate for its metabolic deficiencies, including synthesis of arginine [2], tryptophan [3], purines [4] and cholesterol [5]. *T. gondii* must scavenge these metabolites, along with energy sources like glucose and glutamine, from the host cell [6-8]. As a eukaryote *T. gondii* possesses a complex metabolism with most of the major metabolic pathways complete, including the tricarboxylic acid (TCA) cycle, the pentose phosphate pathway (PPP), gluconeogenesis and glycolysis [6-13]. Other areas of metabolism, including amino acid and nucleotide synthesis, are partially present but rely on host metabolism to provide certain precursors and products [2-4,12,14]. *T. gondii* metabolism also contains unusual pathways for a eukaryote, including a fungal-like shikimate pathway for aromatic amino acid synthesis and an algal-like fatty acid synthesis II pathway [15-17].

Recent metabolic studies have greatly expanded our understanding of *T. gondii* metabolism. The *T. gondii* TCA cycle was found to be essential to growth and a GABA shunt was discovered that could fuel the TCA cycle with glutamine [6]. In no glucose environments, or with glycolysis genetically ablated, *T. gondii* can perform glutaminolysis to provide carbon for gluconeogenesis [7-8]. The fatty acid synthesis II system was shown to synthesize essential long and very long chain fatty acids that could not be provided by host metabolism [18]. A gluconeogenic enzyme, fructose bisphosphatase 2, was found to be constitutively expressed and essential to growth [11]. These and other studies are critical to expanding our understanding of *T. gondii* metabolism; however, the host metabolome has been largely unexplored. For other pathogens, studies that include the host metabolome have led to novel findings, including the discovery of a new metabolic gene in *Plasmodium falciparum* [19].

This current study expands the scope of *T. gondii* infection metabolomics to include the host metabolome. We generated the joint host-parasite metabolome over the course of infection in a human tissue culture infection model, along with matched uninfected control samples. We paired this metabolite analysis with transcriptomics of host and parasite over the same time course of infection. Pairing the joint metabolome with the individual transcriptomes allows us to identify the likely source of metabolic changes based on altered metabolic enzyme transcript abundance in *T. gondii* or the host. We found that the infection metabolome changes over the time course of infection and differs from the uninfected control in several areas. Infection changed amino acid and nucleotide synthesis, glycolysis, the TCA cycle and the PPP. Some changes, such as the TCA cycle, were driven by *T. gondii*, while others, like the PPP, were altered by the host and parasite. Increases in the abundance of sedoheptulose-7-phosphate and sedoheptulose-1,7-bisphosphate led to the discovery of a unique parasite metabolic enzyme, sedoheptulose bisphosphatase, which is not present in mammalian host cells. Further experiments with genetically manipulated parasites showed that *T. gondii* sedoheptulose bisphosphatase energetically drives the synthesis of ribose-5- phosphate. This novel activity gives *T. gondii* a second method for rapidly synthesizing ribose and acts as a metabolic switch to redirect carbon from glycolysis into ribose synthesis. This analysis expands our understanding of host and parasite metabolism during *T. gondii* infection, and highlights a new parasite metabolic enzyme that could be a future drug target.

## Results

### The *T. gondii*-host joint metabolome changes throughout infection

We conducted a temporal mass spectrometry-based analysis of metabolism during *T. gondii* infection using a fibroblast tissue culture infection model. Metabolism was rapidly quenched and metabolites were extracted in triplicate from infected and uninfected dishes at nine time points (1.5, 3, 6, 9, 12, 24, 36, and 48 hours post infection). The metabolome in each sample was quantified using Ultra High-Pressure Liquid Chromatography paired with Mass Spectrometry. We found that *T. gondii* infection dramatically changes metabolism, and that the joint host-parasite metabolome changes over the course of infection. Metabolite abundance was altered for intermediates of central metabolic pathways including glycolysis, the TCA cycle, the PPP, amino acid and nucleotide synthesis (Fig 1).

**Fig 1.**
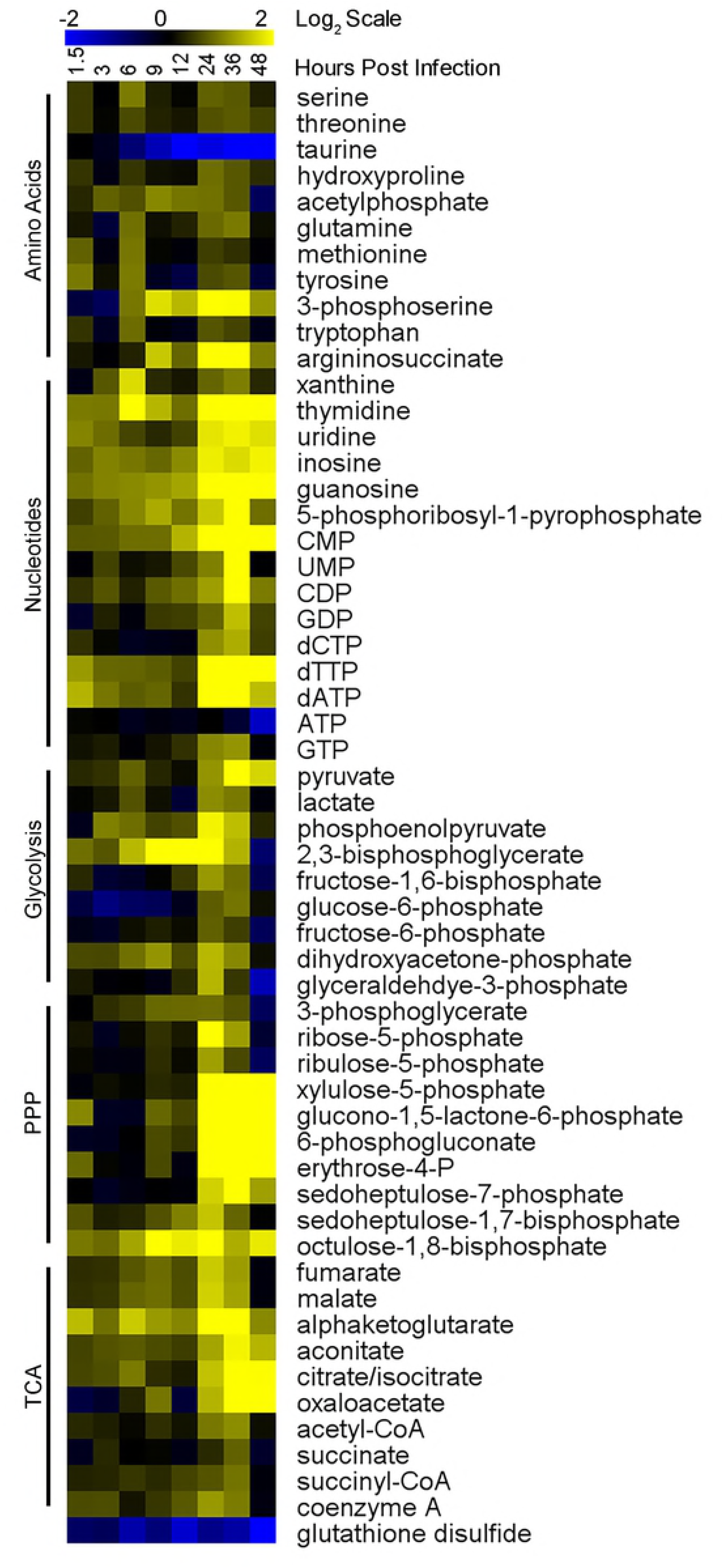
*T. gondii* infection changes the metabolome. Heatmaps show metabolite abundance over 48 hours of *T. gondii* infection. Triplicate infected and uninfected dishes of HFFs were metabolically quenched and metabolites were extracted at 8 times points over time (1.5, 3, 6, 9, 12, 24, 36, and 48 HPI). Metabolomes were quantified using HPLC-MS and metabolites were identified with known standards. Infected sample abundances were averaged and normalized to the average uninfected abundance then log base 2 transformed (Log_2_(Infected Abundance/Uninfected Abundance)) with blue being less abundant and yellow more abundant. Metabolites were sorted by area of metabolism, with significant changes occurring in amino acid and nucleotide synthesis, glycolysis, the TCA cycle, and the PPP.

#### Amino Acid Metabolism

Many amino acids and their biosynthetic intermediates had moderately increased abundance at 24 and 36 hours post infection (HPI) (Fig 1). *T. gondii* is auxotrophic for tryptophan and arginine [2-3], so these amino acids must be scavenged from the host cell. Tryptophan abundance increased over the infection time course. Likewise, argininosuccinate, the last metabolite in arginine synthesis before arginine, dramatically increased abundance in infected cells. The infected host transcriptome compared to the uninfected control indicates possible explanations for these metabolomic shifts. Humans lack the ability to synthesize tryptophan and LAT1 is the transporter responsible for tryptophan uptake. Transcription of LAT1 increases during infection, particularly from 6 to 48 HPI when expression is 5-fold higher than the uninfected control (Table 1). Arginino Succinate Synthase 1, which catalyzes the rate limiting step in arginine synthesis, was highly expressed throughout infection, including a period from 9 to 24 HPI when expression was roughly double that of the uninfected cells (Table 1). 3-phosphoserine abundance also increased during infection, which likely indicates an increase in serine synthesis. The rate limiting step in serine synthesis is catalyzed by 3-phosphoglycerate dehydrogenase (PHGDH), which is highly expressed in host and parasite, indicating they likely both contribute to the increase in serine. In infected host cells, PHGDH expression is over 3.5 fold upregulated from 12 to 36 HPI, while *T. gondii* PHGDH is expressed at a consistently high level, over 150 Fragments Per Kilobase of transcript per Million mapped (FPKM), from 3 through 48 HPI (Table 2).

**Table 1.** Host mRNA abundance of genes of interest. mRNA abundance of host genes of interest in infected cells normalized to uninfected cell expression. RNA samples were isolated from infected dishes paired to infected metabolomics dishes and from an uninfected control dish. RNAseq was performed to generate the transcriptome and infected cell expression was normalized to the uninfected baseline to determine how expression was changed by *T. gondii* infection. Infection increased expression of multiple host metabolic genes in central pathways including glycolysis, the PPP, nucleotide metabolism, and amino acid synthesis and transport. Large amino acid transporter 1 (LAT1), argininosuccinate synthase 1 (ASS1), phosphoglycerate dehydrogenase (PHGDH), methylenetetrahydrofolate dehydrogenase (MTHFD1), 5- phosphoribosyl-1-pyrophosphate synthase (PRS1), hexose kinase (HK), pyruvate kinase (PK), lactate dehydrogenase A (LDHA), bisphosphoglycerate mutase (BPGM), aconitase (ACN), isocitrate dehydrogenase (IDH), alpha-ketoglutarate dehydrogenase (A-KGDH), succinate reductase (SCR), fumarase (FUM), malate dehydrogenase (MDH), citrate synthase (CS), succinyl-CoA synthetase (SCS), glucose-6-phosphate dehydrogenase (G6PD), 6-phosphogluconolactonase (6PGL), 6-phosphogluconate dehydrogenase (6PGD), ribose-5-phosphate isomerase (R5PI), ribulose-5-phosphate epimerase (Ru5PE), transaldolase (TA), transketolase (TK).

| Human Genes of Interest (Infection FPKM normalized to uninfected) |  |  |  |  |  |  |  |
| --- | --- | --- | --- | --- | --- | --- | --- |
| Gene | 1.5 | 6 | 9 | 12 | 24 | 36 | 48 |
| LAT1 | 5.49 | 11.5 | 5.89 | 11.8 | 6.31 | 4.89 | 51.7 |
| ASS1 | 1.27 | 1.92 | 1.96 | 2.39 | 2.29 | 1.48 | 0.35 |
| PHGDH | 0.80 | 2.30 | 1.14 | 3.55 | 5.46 | 4.90 | 1.40 |
| MTHFD1 | 0.77 | 1.47 | 1.30 | 1.59 | 2.06 | 2.58 | 2.05 |
| PRS1 | 2.27 | 2.26 | 2.18 | 2.21 | 3.12 | 3.94 | 2.64 |
| MYC | 1.99 | 6.07 | 4.85 | 5.23 | 3.04 | 1.76 | 4.30 |
| HK | 1.36 | 3.20 | 2.85 | 2.65 | 1.50 | 1.81 | 2.88 |
| PK | 1.25 | 1.44 | 1.30 | 1.44 | 1.33 | 1.51 | 1.12 |
| LDHA | 1.79 | 2.85 | 2.53 | 2.22 | 2.34 | 2.51 | 0.91 |
| BPGM | 1.50 | 1.36 | 1.56 | 0.71 | 0.52 | 0.44 | 0.18 |
| ACN | 1.08 | 0.92 | 0.95 | 0.89 | 0.58 | 0.66 | 0.56 |
| IDH | 1.05 | 0.97 | 1.12 | 1.03 | 1.03 | 1.18 | 0.39 |
| A-KGDH | 0.79 | 0.96 | 0.99 | 0.83 | 0.64 | 0.50 | 1.48 |
| SCR | 0.99 | 0.68 | 0.78 | 0.79 | 0.90 | 1.19 | 0.22 |
| FUM | 0.69 | 0.62 | 0.63 | 0.57 | 0.65 | 0.59 | 0.20 |
| MDH | 0.93 | 0.91 | 0.93 | 0.94 | 1.02 | 0.98 | 0.34 |
| CS | 1.01 | 2.65 | 1.50 | 2.18 | 1.28 | 1.26 | 4.44 |
| SCS | 0.70 | 0.49 | 0.42 | 0.57 | 0.56 | 0.31 | 0.48 |
| G6PD | 1.08 | 1.35 | 1.73 | 1.63 | 1.28 | 1.15 | 0.42 |
| 6PGL | 0.78 | 0.66 | 0.62 | 0.75 | 0.95 | 0.83 | 0.18 |
| 6PGD | 0.86 | 1.55 | 1.33 | 1.68 | 2.31 | 3.25 | 1.55 |
| R5PI | 1.29 | 2.60 | 2.54 | 2.88 | 2.50 | 2.05 | 1.79 |
| Ru5PE | 0.58 | 0.95 | 0.57 | 1.13 | 0.76 | 1.11 | 1.10 |
| TA | 1.03 | 1.32 | 1.42 | 1.62 | 2.46 | 2.19 | 0.35 |
| TK | 0.59 | 0.65 | 0.61 | 0.52 | 0.59 | 0.68 | 0.32 |

**Table 2.** *T. gondii* mRNA abundance of genes of interest. mRNA abundance of *T. gondii* genes of interest in raw FPKM values. RNA samples were isolated from infected dishes paired to infected metabolomics dishes and RNAseq was performed to generate the transcriptomes. There is no baseline to normalize *T. gondii* expression to, as the parasite cell cycle is constantly shifting. Genes throughout central metabolism are expressed at a high level throughout the parasite lytic cycle. Phosphoglycerate dehydrogenase (PHGDH), hypoxanthine phosphoribosyltransferase (HXGPRT), adenosine kinase (AK), 5-phosphoribosyl-1-pyrophosphate synthase (PRS1), hexose kinase (HK), pyruvate kinase (PK), lactate dehydrogenase 1 (LDH1), aconitase (ACN), alpha-ketoglutarate dehydrogenase (A-KGDH), succinate reductase (SCR), fumarase (FUM), isocitrate dehydrogenase 2 (IDH – TCA), isocitrate dehydrogenase 1 (IDH-APICO), malate dehydrogenase (MDH), citrate synthase (CS), succinyl-CoA synthetase (SCS), glucose-6-phosphate dehydrogenase putative 1 (G6PD 1), glucose-6-phosphaye dehydrogenase putative 2 (G6PD 2), 6-phosphogluconate dehydrogenase putative 1 (6PGD 1), 6-phosphogluconate dehydrogenase putative 2 (6PGD 2), ribose-5-phosphate isomerase (R5PI), ribulose-5-phosphate epimerase (Ru5PE), transaldolase (TA), transketolase (TK), sedoheptulose bisphosphatase (SBPASE).

| *Toxoplasma* Genes of Interest (Unprocessed FPKM values) |  |  |  |  |  |  |  |
| --- | --- | --- | --- | --- | --- | --- | --- |
| Gene | 1.5 | 6 | 9 | 12 | 24 | 36 | 48 |
| PHGDH | 131 | 221 | 228 | 217 | 215 | 190 | 157 |
| HXGPRT | 7.85 | 23.3 | 21.2 | 17.7 | 20.9 | 8.72 | 56.6 |
| AK | 21.3 | 52.9 | 55.1 | 48.3 | 51.0 | 40.9 | 16.9 |
| PRS1 | 28.2 | 38.5 | 34.2 | 28.8 | 23.5 | 16.6 | 35.9 |
| HK | 42.4 | 72.7 | 79.9 | 68.5 | 70.3 | 60.7 | 59.8 |
| PK | 50.0 | 116 | 95.7 | 96.8 | 93.3 | 79.3 | 116 |
| LDH1 | 351 | 361 | 394 | 337 | 349 | 310 | 614 |
| ACN | 27.9 | 33.0 | 24.1 | 37.0 | 26.9 | 21.5 | 52.8 |
| A-KGDH | 18.3 | 20.9 | 18.5 | 25.2 | 19.9 | 14.8 | 32.4 |
| SCR | 27.8 | 38.6 | 28.2 | 37.7 | 28.9 | 21.6 | 39.5 |
| FUM | 8.63 | 15.7 | 18.4 | 16.2 | 13.8 | 9.28 | 13.0 |
| IDH-TCA | 56.1 | 110 | 115 | 112 | 106 | 94.0 | 42.3 |
| IDH-APICO | 3.38 | 44.2 | 23.9 | 45.0 | 43.7 | 38.9 | 13.3 |
| MDH | 79.4 | 259 | 248 | 280 | 309 | 218 | 52.7 |
| CS | 24.1 | 39.0 | 39.5 | 44.6 | 42.4 | 38.2 | 26.8 |
| SCS | 75.8 | 103 | 114 | 116 | 119 | 93.5 | 61.0 |
| G6PD 1 | 39.5 | 41.9 | 27.9 | 40.1 | 21.1 | 21.3 | 127 |
| G6PD 2 | 710 | 435 | 434 | 364 | 292 | 363 | 466 |
| 6PGD 1 | 0.00 | 0.00 | 0.00 | 0.08 | 0.00 | 0.17 | 0.09 |
| 6PGD 2 | 4.69 | 22.3 | 34.6 | 19.5 | 25.2 | 23.3 | 13.9 |
| R5PI | 16.8 | 31.1 | 31.0 | 35.4 | 23.5 | 22.6 | 16.9 |
| Ru5PE | 10.0 | 14.4 | 21.4 | 12.3 | 15.2 | 14.7 | 15.6 |
| TA | 77.6 | 122 | 134 | 103 | 112 | 126 | 47.6 |
| TK | 6.96 | 40.3 | 38.1 | 39.0 | 35.8 | 22.0 | 27.6 |
| SBPASE | 57.1 | 98.8 | 118 | 91.3 | 105 | 112 | 38.5 |

#### Nucleotide Synthesis

A wide range of metabolites involved in nucleotide synthesis dramatically increased abundance in infected cells, with most changes occurring in the 24-48 HPI range (Fig 1). These increased metabolites include guanosine, inosine, uridine, deoxynucleoside thymidine, xanthine and 5-phosphoribosyl-1-pyrophosphate (PRPP), which is an essential metabolite in purine and pyrimidine synthesis and salvage. Additionally, multiple nucleotides (CMP, UMP, CDP, GDP, and GTP) and deoxy nucleotides (dCTP, dTTP, and dATP) increased abundance during infection, while only ATP had a slightly decreased abundance late in infection. In infected cells, the important purine synthetic enzyme tetrahydrofolate synthase (MTHFD1) was upregulated compared to uninfected cells from 6 to 48 HPI (Table 1). PRPP Synthase (PRS1), which is essential for purine and pyrimidine synthesis, was upregulated in infected host cells throughout infection, with a peak of 4 fold increased expression at 36 HPI (Table 1). Transcription of the regulatory protein Myc, which activates nucleotide synthesis, was increased at every time point in infected cells, reaching 6 fold increased transcription over uninfected cells at 6 HPI (Table 1). This result was expected given previous work demonstrating *T. gondii* infection induces increased Myc expression and activity [20]. Two parasite enzymes which are essential to nucleotide salvage, hypoxanthine-guanine phosphoribosyl transferase and adenosine kinase, were consistently expressed throughout infection, as was *T. gondii* PRS1 which creates the PRPP needed for nucleotide synthesis and salvage (Table 2).

#### Glycolysis

Glycolytic intermediates were generally more abundant in the infection metabolome, apart from 48 HPI when many were depleted. This result is understandable given previous work demonstrating a highly active *T. gondii* glycolytic pathway and the likely energetic strain that infection puts on host cells [8,11,21]. Host hexose kinase (HK) and pyruvate kinase (PK) were transcriptionally upregulated in infected cells at every time point during infection (Table 1) and were highly transcribed throughout infection in *T. gondii* with FPKM values regularly at or above 50 (Table 2). Unexpectedly 2,3-bisphosphoglycerate (2,3-BPG) and lactate were also increased in abundance in the infection metabolome. Lactate dehydrogenase (LDH) was transcriptionally upregulated in infected host cells and highly transcribed throughout infection in *T. gondii*, meaning that both likely contributed to the increase in lactate (Tables 1 and 2). 2,3-BPG is synthesized in the rapport-leubering shunt, which decreases ATP synthesis from glycolysis by 50% [22]. Bisphosphoglycerate mutase (BPGM), the main source of 2,3-BPG, is transcriptionally upregulated in infected host cells during the first twelve hours of infection (Table 1). *T. gondii* BPGM has yet to be identified, which makes it difficult to estimate the parasite contribution to 2,3-BPG synthesis.

### Increase in TCA cycle metabolites is likely driven by *T. gondii* and not the host

*T. gondii* infection of fibroblasts increases the abundance of multiple TCA cycle intermediates, including citrate/isocitrate, oxaloacetate, aconitate and α-ketoglutarate (Fig 2). Changes in abundance range from 2-8 fold increases over uninfected cells from 24 to 48 HPI. Malate, fumarate, succinyl-CoA and succinate were also increased in abundance 1.5 to 2 fold over uninfected cells from 24 to 48 HPI. Two key metabolites, acetyl-CoA and Coenzyme A, which are essential to fueling the TCA cycle, are similarly more abundant during late *T. gondii* infection. Taken together the metabolomic data demonstrates that the *T. gondii*-Host infection metabolome has a highly active TCA cycle, particularly late during infection.

**Fig 2.**
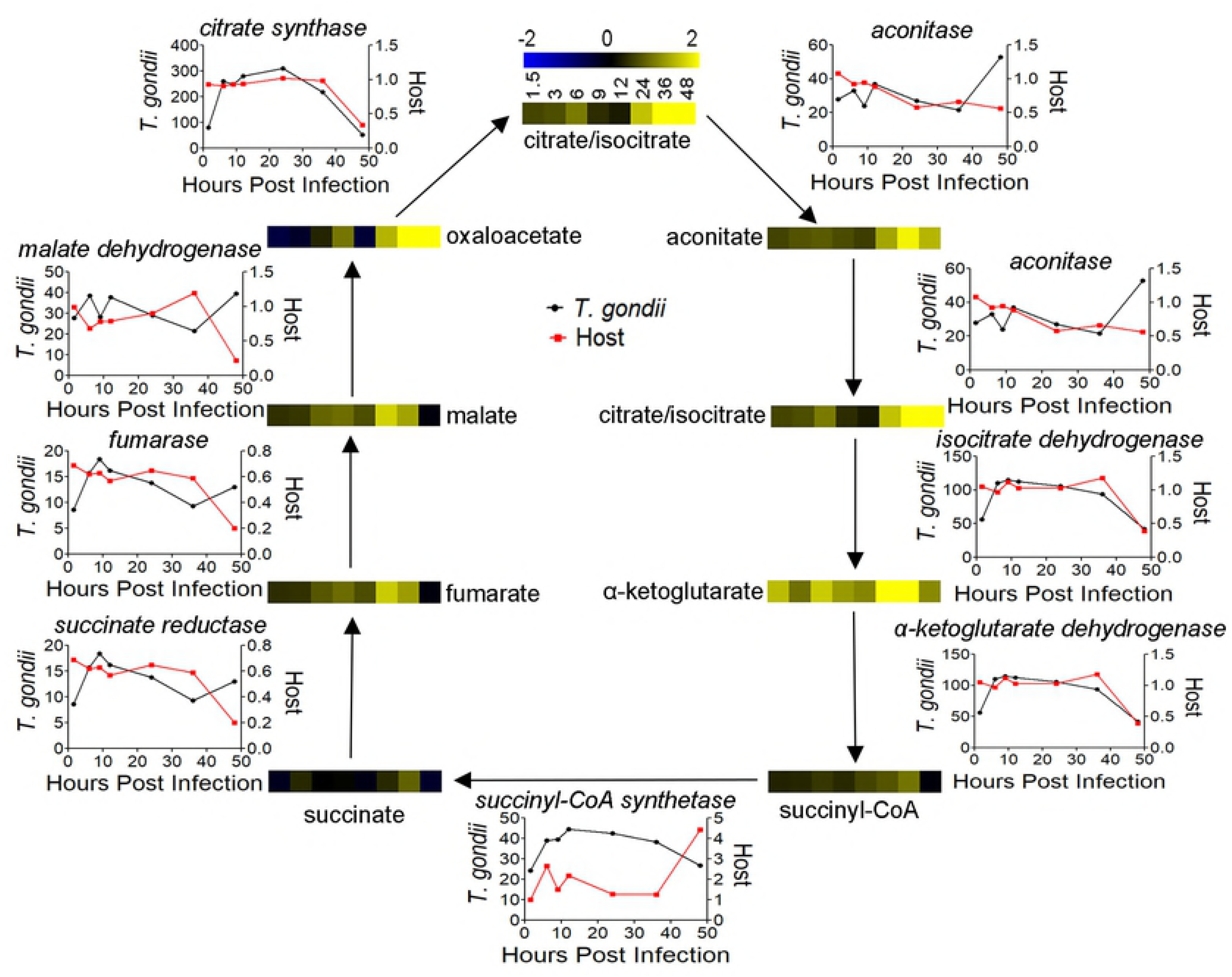
*T. gondii* infection TCA cycle metabolome and transcriptome. Heatmaps show abundance of each TCA cycle intermediate over 48 hours with blue being less abundant and yellow more abundant. The line graphs represent mRNA abundance for each host (red) and *T. gondii* (black) TCA cycle enzymes with names italicized. Host expression is normalized to uninfected transcriptome expression, *T. gondii* expression is shown in raw FPKM values. Large increases in metabolite abundance during late infection paired with consistent expression of *T. gondii* TCA enzymes indicate parasite driven increases in TCA intermediate abundance.

Our *T. gondii* transcriptome supports previous findings, with all enzymes in the TCA cycle expressed throughout infection (Table 2). Critically, the enzyme that catalyzes the rate limiting step of the TCA cycle, Isocitrate dehydrogenase, is highly expressed at roughly 100 FPKM from 6 to 36 HPI. The enzyme which is responsible for generating acetyl-CoA to fuel the TCA cycle, branched-chain α-ketoacid dehydrogenase (BCKDH), is also expressed at a high level throughout infection. There are secondary forms of two TCA cycle enzymes, aconitase and isocitrate dehydrogenase, which are localized to the apicoplast where they produce reducing equivalents for biosynthetic pathways [23]. While these enzymes are not involved in the mitochondrial TCA cycle, they share intermediates and contribute to the observed TCA cycle phenotype. The apicoplast-aconitase originates from the same gene as the mitochondrial aconitase but is localized to both organelles through an unusual signal sequence [23]. The apicoplast isocitrate dehydrogenase is highly expressed from 6 to 48 HPI and likely plays a role in the increase in abundance of citrate/isocitrate and α-ketoglutarate.

While the *T. gondii* TCA cycle is known to be active during infection, the host cycle has not been as heavily studied. Transcription of most host TCA cycle genes does not increase during infection (Table 1). Host succinate dehydrogenase, succinyl-CoA synthetase, and fumarase are less abundant in infected cells throughout, while pyruvate dehydrogenase, α-ketoglutarate dehydrogenase, aconitase, malate dehydrogenase, and isocitrate dehydrogenase are at or below the expression of uninfected control cells. The only enzyme with increased abundance during infection was citrate synthase, which appears to be moderately increased in expression. Taken together these two transcriptomes indicate the origins of the increases in the infection metabolome TCA cycle. The increase in abundance of TCA cycle intermediates is driven by the highly active *T. gondii* TCA cycle and replication of the parasite, while the host contribution to the TCA cycle appears to be minimal, with key enzymes either repressed or barely transcribed during infection.

### The Pentose Phosphate Pathway (PPP) is activated in *T. gondii* and the host

The abundance of key intermediates in the oxidative and non-oxidative PPP are dramatically increased in the infection metabolome. Glucono-1,5-lactone-6-phosphate and gluconate-6-phosphate in the oxidative pathway are increased in abundance from 2.5 to 4.9 fold over uninfected cells from 24 to 48 HPI (Fig 3). A similar pattern was found in xylulose-5-phosphate abundance, which was 2.8 to 3.9 fold more abundant from 24 to 48 HPI. Sedoheptulose-7-phosphate was 1.6 to 2.2 fold more abundant from 24 to 48 HPI, with similar changes occurring in sedoheptulose-1,7-bisphosphate. Octulose-1,8-bisphosphate followed the same pattern as sedoheptulose-1,7- bisphosphate and although the abundance was too low to be significant. Octulose-8- phosphate followed a similar pattern to sedoheptulose-7-phosphate (Fig 1). Erythrose-4-phosphate abundance was increased during late infection despite being undetectable in multiple uninfected samples. Glyceraldehyde-3-phosphate was moderately more abundant in infected samples but to a much lesser degree than other PPP intermediates and shifts in abundance were not limited to late infection. Given the dual role of glyceraldehyde-3-phosphate as an intermediate in glycolysis and the PPP, it is understandable that shifts in its abundance do not match up perfectly with other metabolites in either pathway.

**Fig 3.**
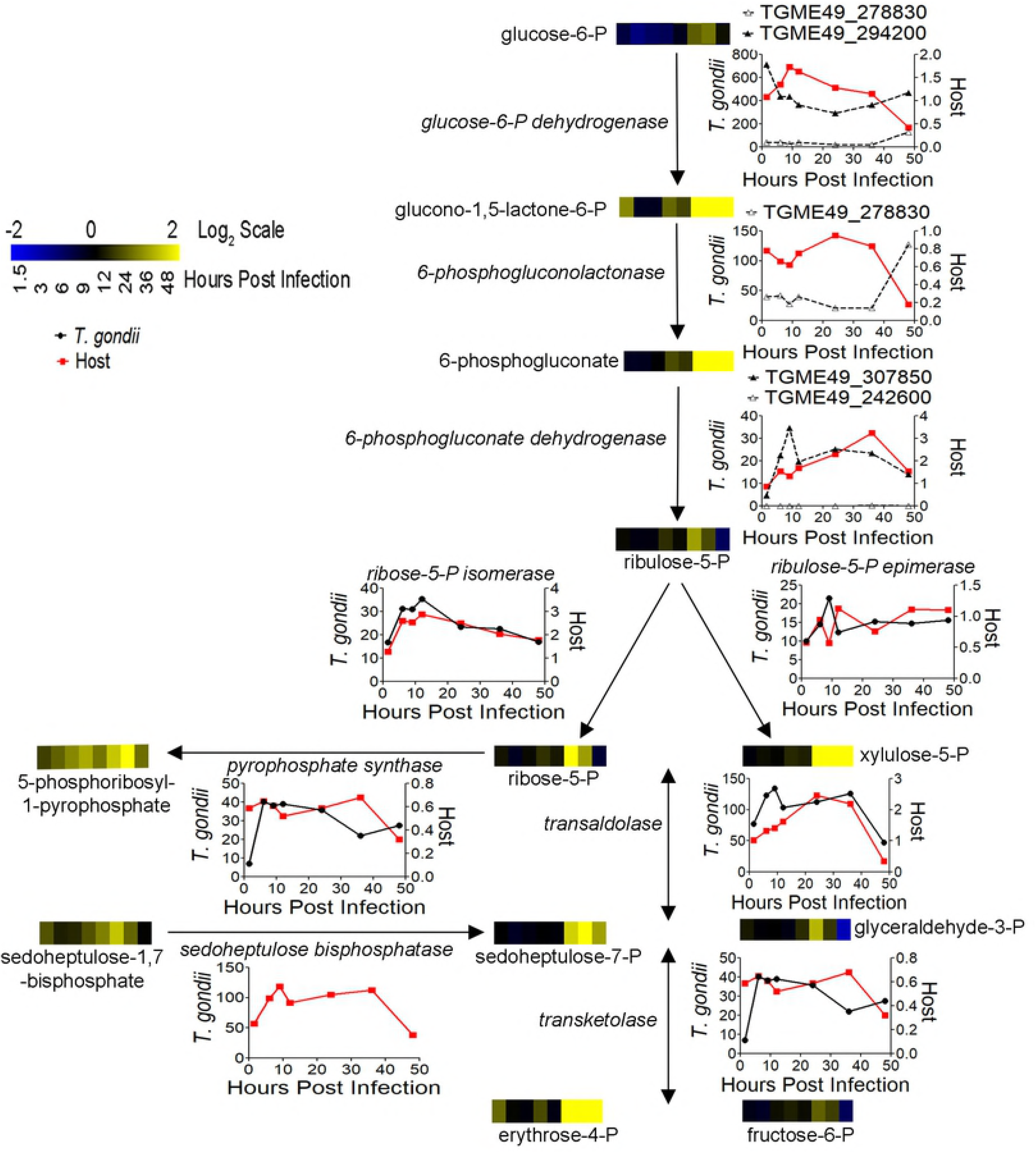
*T. gondii* infection PPP metabolome and transcriptome. Heatmaps show abundance of each PPP intermediate over 48 hours. The graphs represent mRNA abundance for each host (red) and *T. gondii* (black) PPP cycle enzyme with enzyme titles italicized. Host expression is normalized to uninfected transcriptome expression, *T. gondii* expression is in raw FPKM values. Putative *T. gondii* genes in the oxidative PPP are noted with dashed lines. Large increases in metabolite abundance during late infection paired with consistent expression of host and *T. gondii* PPP enzymes indicate both host and parasite drive increases in PPP intermediate abundance.

The host transcriptome supports an active oxidative and non-oxidative PPP (Table 1). The rate limiting enzyme of the oxidative pathway, glucose-6-phosphate dehydrogenase, is expressed at or slightly above uninfected levels for the majority of infection, 6 to 36 HPI. Messenger RNA abundance of 6-phosphogluconolactone dehydrogenase, which catalyzes the third reaction in the oxidative PPP, is higher in infected cells from 6 to 48 HPI, peaking at 3.3 fold at 36 HPI. Both steps are responsible for generating NADPH and their expression strongly indicates an activated host oxidative PPP. The two enzymes of the non-oxidative PPP, transketolase and transaldolase, have minor regulation but are expressed throughout infection.

Transcriptomic analysis of the *T. gondii* PPP is complicated because the genes of the oxidative PPP are uncharacterized. Based on the current genome annotation there are two putative glucose-6-phosphate dehydrogenases and two putative 6-phosphogluconate dehydrogenases. One of the putative glucose-6-phosphate dehydrogenase genes also has putative phosphogluconolactonase activity. This activity would be in agreement with recent findings that *Plasmodium* possesses a bifunctional enzyme responsible for the first two steps of the oxidative PPP [24]. The putative bifunctional glucose-6-phosphate dehydrogenase-phosphogluconolactonase was expressed throughout infection, with FPKM values in the 20 to 40 range, while the glucose-6-phosphate dehydrogenase without putative secondary activity was highly expressed with FPKM’s over 300 for almost all of infection (Table 2). In contrast, only one of the 6-phosphogluconate dehydrogenases was expressed throughout infection, with FPKMs in the 20-30 range, while the other had no expression at multiple time points. Given that at least one of each of the putative oxidative PPP genes are expressed throughout infection we believe that the *T. gondii* oxidative PPP is active during infection. In the non-oxidative PPP *T. gondii* transketolase is expressed at 20 to 40 FPKM from 6 to 48 HPI while transaldolase is heavily expressed throughout infection with FPKM values over 100 from 6 to 36 HPI. Taken together, the host and parasite transcriptomes highlight that the increase in PPP metabolite abundance in the infection metabolome is likely due to increased activity in the oxidative and non-oxidative halves of the PPP in both *T. gondii* and its host cell.

### *T. gondii* possesses Sedoheptulose Bisphosphatase activity

Increased abundance in sedoheptulose-7-phosphate (S7P) and sedoheptulose-1,7-bisphosphate (SBP), which repeated in octulose-8-phosphate (O8P) and octulose-1,8-bisphosphate (OBP), led us to investigate whether these metabolites were linked. Sedoheptulose-1,7-bisphosphatase (SBPase) in *Saccharomyces cerevisiae* is known to generate S7P and O8P from SBP and OBP respectively [25]. SBPase activity does not exist in warm-blooded animals; however, *Toxoplasma* has a putative SBPase (TGME49_235700) and genetic analysis has identified a likely SBPase in the protozoan parasite *Trypanosoma brucei* [26].

SBPase activity can be assayed for using Glucose-6-^13^C isotope labeling. In the absence of SBPase, S7P will only be present in the unlabeled or +1 labeled forms, while cells with SBPase activity will have +2 and +3 labeled forms [25]. Triplicate infected and uninfected dishes were grown for 24 hours with Glucose-6-^13^C media. Uninfected S7P was 41% unlabeled and 59% +1 labeled, exhibiting no SBPase activity (Figs 4A and S1). Infected S7P was 30% unlabeled, 46% +1 labeled, 21% +2 labeled, and 3% +3 labeled, indicating significant SBPase activity (Fig 4A). The absence of +2 labeled and +3 labeled S7P in the uninfected cells was not due to a lack of +2 labeled SBP (Fig 4B). Uninfected cells had 30% +2 labeled, and 2.8% +3 labeled SBP, demonstrating that uninfected cells cannot convert SBP into S7P (Fig 4B).

**Fig 4.**
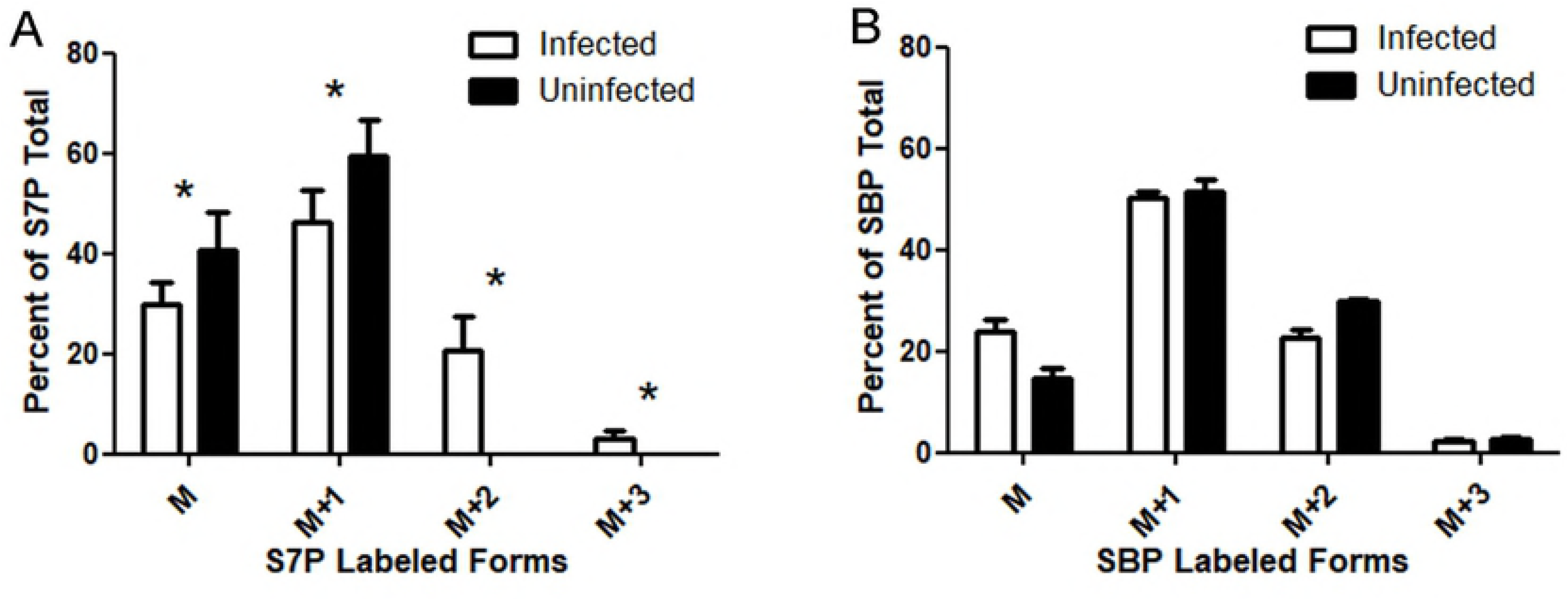
*T. gondii* possesses SBPase activity. (A) Mass (M) of S7P as a percentage of the total in infected (white bars) and uninfected (black bars) cells. M+1 is S7P containing one ^13^C, M+2 is S7P containing two ^13^C, and M+3 is S7P containing three ^13^C. Asterisks mark significant differences (p>0.05, two tailed t-test) between infected and uninfected samples. Error bars are a 95% confidence interval. (B) Mass (M) of SBP as a percentage of the total in infected (white bars) and uninfected (black bars) cells. M+1 is SBP containing one 13C, M+2 is SBP containing two 13C, and M+3 is SBP containing three 13C. Error bars are a 95% confidence interval.

To estimate flux through SBPase, we divided the portion of +2 and +3 labeled S7P by the +2 and +3 labeled portion of SBP because the only source of +2 or +3 labeled S7P is the dephosphorylation of +2 or +3 labeled SBP. These calculations estimated 91% flux through SBPase in *T. gondii* infected cells, indicating that the majority of observed S7P in infected cells is from SBPase activity. These results highlight that *T. gondii* possesses a novel central metabolic pathway catalyzed by a highly active enzyme which humans lack.

### Expression of *T. gondii* SBPase in HeLa cells shows SBPase activity

To confirm the identity of the putative *T. gondii* SBPase (TGME49_235700) a lentiviral expression system was used to stably incorporate the gene into the genome of HeLa cells. As a control, a separate population of HeLa cells were transduced with Blue Fluorescent Protein (BFP). Triplicate 70% confluent 60 mm dishes were grown for 3 hours with Glucose-6-^13^C media (Fig 5). In BFP expressing cells, S7P was 15% unlabeled, 85% +1 labeled. In SBPase expressing cells S7P was 22% unlabeled, 72% +1 labeled and 6% +2 labeled. Fluorescence microscopy of the drug selected populations confirmed successful transduction and expression of BFP and no cross-contamination of lentiviruses during transduction (Fig S2). qPCR analysis confirmed that SBPase cells were expressing SBPase at 3.2% of ActB expression and BFP cells had no measurable SBPase expression.

**Fig 5.**
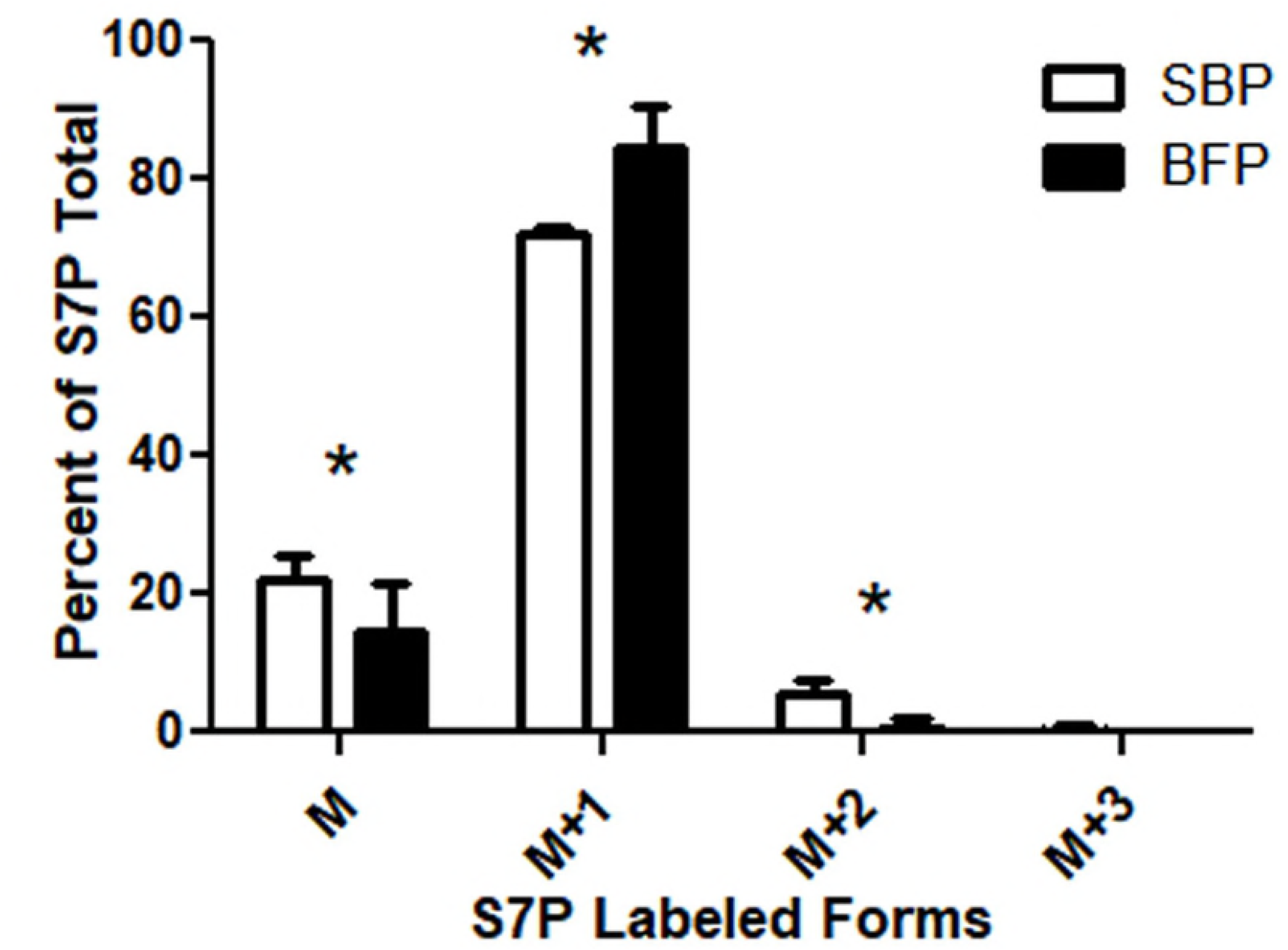
Expression of *T. gondii* SBPase in HeLa results in SBPase activity. Average S7P isotope composition for HeLa cells expressing SBPase (SBP, white bars) or BFP (black bars) as a percentage (M= parental unlabeled, M+1 = mono ^13^C labeled, M+2 = double ^13^C labeled). The significant pool of M+2 S7P in SBPase expressing samples confirms TGME49_235700 codes for SBPase while a lack of M+2 S7P in BFP expressing samples confirms that host cells lack this enzyme. Asterisks mark significant differences (p>0.05, two tailed t-test) between SBPase and BFP expression samples. Error bars are a 95% confidence interval.

### Overexpression of *T. gondii* SBPase alters infection metabolome

SBPase was expressed from the constitutive *T. gondii* α-tubulin promoter randomly electroporated into the *T. gondii* genome. Overexpression clones (SBPOE) were assayed for with southern blotting and two unique clones, SBPOE1 and SBPOE2, were isolated (Fig S3). A limited metabolomics time course was carried out using Wild Type (WT) ME49, SBPOE1 and SBPOE2, with samples taken at 9, 12, 24, and 36 HPI. The metabolomes of each SBPOE clone were normalized to the WT. While the two clones displayed similar metabolomic shifts the amplitude of these changes was much greater in SBPOE2 than SBPOE1 (Fig 6). qPCR analysis found WT SBPase expression was 57.2% ± 4.3 of Tub1a, while SBPOE1/Tub1a was 179% ± 20.5 and SBPOE2 /Tub1a was 256% ± 62.6. Thus, the qPCR data confirmed that SBPOE2 expresses SBPase at a higher level than SBPOE1 although both express SBPase at above WT levels. S7P was 2.1 and 3 fold more abundant at 9 HPI in SBPOE6 and SBPOE2 infected populations, respectively. Multiple nucleotide metabolites increased during SBPOE1 and SBPOE2 infection over WT levels including xanthine, hypoxanthine, UDP, ADP, ATP, inosine, and allantoin. Lactate and fructose-1,6-bisphosphate abundance was also moderately increased in SBPOE1 and SBPOE2. These results indicate that SBPase plays a similar role in *T. gondii* metabolism, energetically driving the synthesis of S7P which drives the synthesis of ribose and increases nucleotide synthesis, with extra S7P flowing into glycolysis.

**Fig 6.**
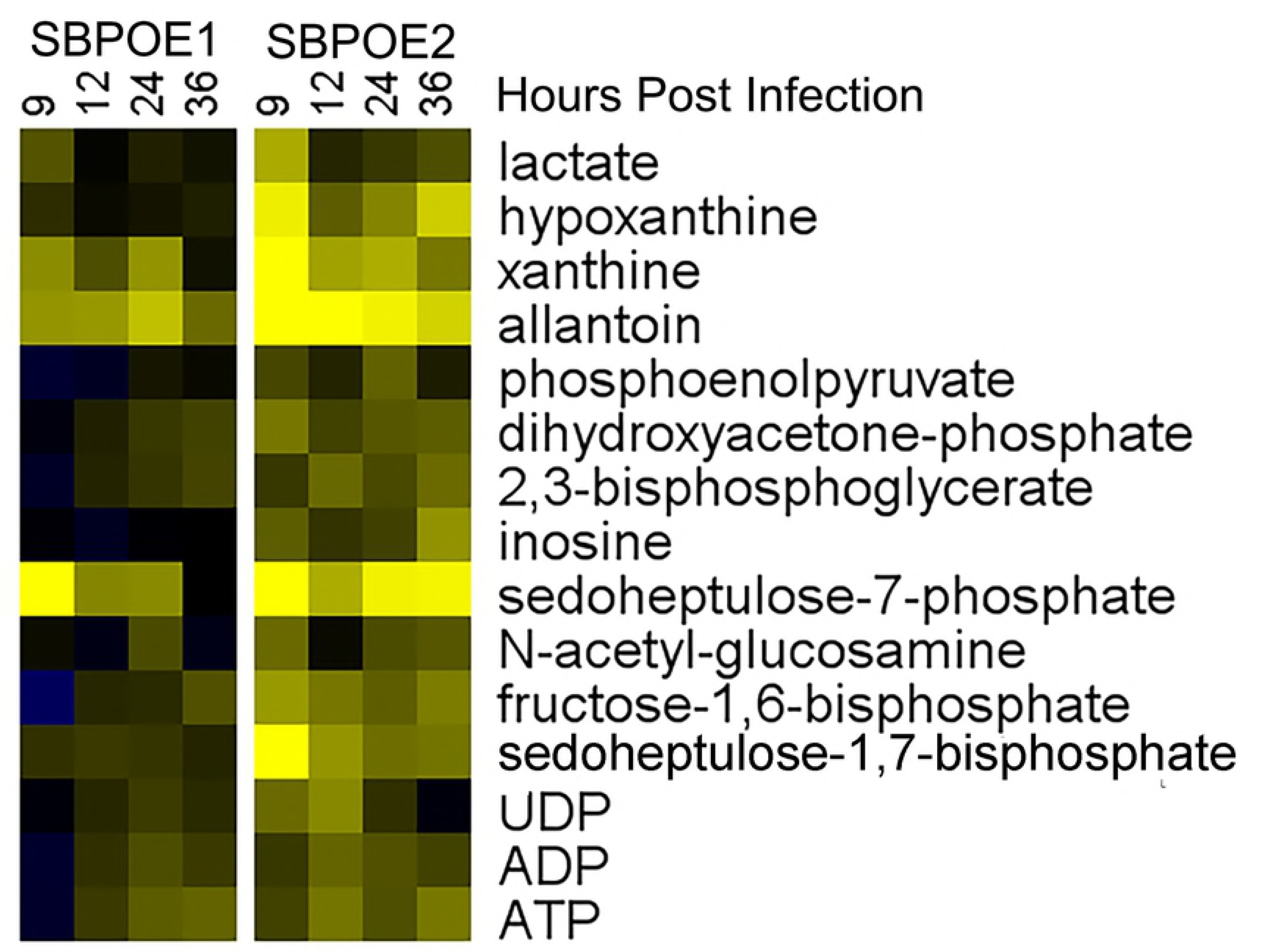
Overexpression of *T. gondii* SBPase Changes Infection Metabolome. Heatmaps show metabolite abundances from *T. gondii* infection time course of 9, 12, 24 and 36 HPI with the same scale of previous heatmaps. Duplicate samples of SBPOE1 (left) or SBPOE2 (right) infection were measured by LC-MS for metabolites. Metabolite abundances were averaged and normalized to the average duplicate WT abundance then log base 2 transformed (Log_2_(SBPOE Abundance/WT Abundance)). Metabolites were sorted by area of metabolism and selected based on known areas of interest. SBPOE1 and SBPOE2 differ from WT in the same areas of metabolism although changes in SBPOE2 metabolite abundance are of greater amplitude than SBPOE1.

### Overexpression of *T. gondii* SBPase increases S7P and flux through the non-oxidative PPP

We used isotopic labeling to determine if SBPOE1 and SBPOE2 had increased flux through SBPase and the non-oxidative PPP. We repeated the SBPase labeling assay in quadruplicate; however, we did not observe an increase in flux. S7P isotopic labeling was indistinguishable between the SBPOE1, SBPOE2, and WT populations. To assay flux through the PPP we labeled SBPOE1, SBPOE2, and WT infected cells in quadruplicate with glucose-1,2-^13^C. The oxidative PPP removes the first carbon, resulting in a +1 labeled form of ribose-5-phosphate, while the non-oxidative PPP does not remove any carbons which gives a +2 pool of ribose-5-phosphate. Ribose-5- phosphate in each sample was heavily C13 labeled (Fig S4). The ratio of +1 to +2 labeled ribose-5-phosphate approximates the ratio of oxidative to non-oxidative PPP flux. SBPOE1 (2.23) and SBPOE2 (2.08) had significantly (P>0.001) lower ratios than WT (3.15) (Fig 7), indicating that flux through the non-oxidative PPP is increased during infection by *T. gondii* overexpressing SBPase.

**Fig 7.**
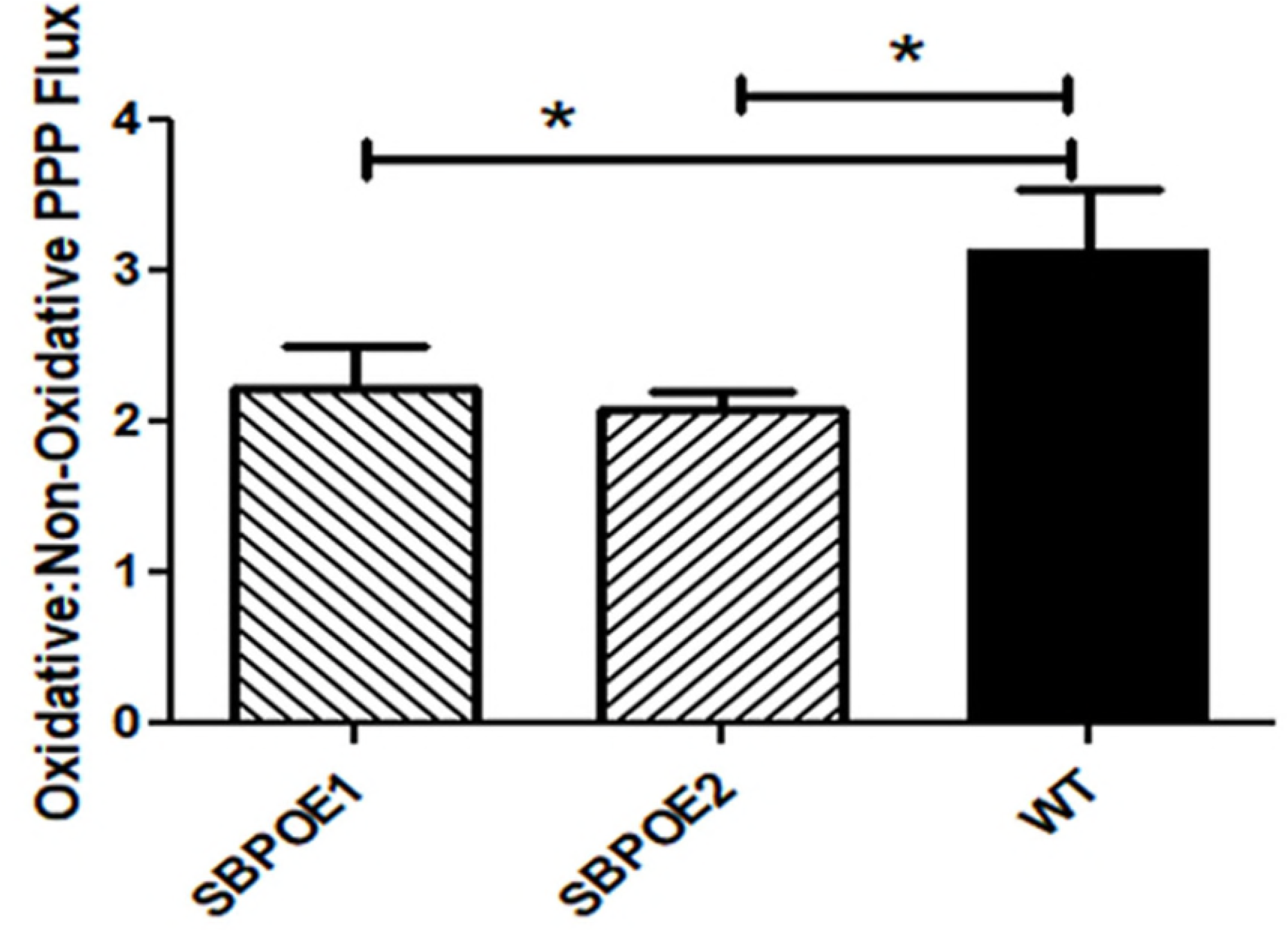
Overexpression of *T. gondii* SBPase increases non-oxidative PPP flux. SBPOE1, SBPOE2, and WT infected cells were grown in quadruplicate with glucose-1,2-13C for 24 hours. Ribose-5-phosphate isotope composition was determined and the average ratio of +1 to +2 labeled forms was calculated for SBPOE1, SBPOE2, and WT. SBPOE1 and SBPOE2 have significantly lower +1:+2 ratios of ribose-5-phosphate which demonstrates a greater relative level of non-oxidative PPP flux. Asterisks mark significant differences (P>0.001, one way ANOVA with Bonferroni’s correction for multiple comparisons) between WT and SBPOE1 or SBPOE2 infected samples. Error bars are a 95% confidence interval.

## Discussion

Recent studies of *T. gondii* metabolism have expanded our understanding of fatty acid synthesis, glycolysis, and the TCA cycle. These studies have found new areas of *T. gondii* metabolism, such as the synthesis of unique fatty acids important for replication [18] and the presence of a GABA shunt in the TCA cycle [6]. Recent *Plasmodium falciparum* metabolomic research included host metabolism, which led to the discovery of a parasite gene that likely contributes to human malarial hypoargininemia [19]. Our study widens the scope of *T. gondii* metabolomics research to include the host metabolism changes after *T. gondii* infection and pairs this dual metabolome with transcriptomic analysis to identify the source of metabolic shifts. We show that *T. gondii* infection changes amino acid synthesis, nucleotide metabolism, glycolysis, the TCA cycle and the PPP. TCA cycle metabolites were likely synthesized primarily by *T. gondii* while increased PPP were driven by the host and parasite. Changes in the PPP led to the discovery of a novel parasite enzyme, sedoheptulose bisphosphatase, which drives ribose-5-phosphate production through the non-oxidative PPP and is not present in host cells.

Several metabolites decreased in abundance during infection, including glutathione disulfide and taurine. Glutathione disulfide depletion may be due to the reduction of free radicals produced by oxidative metabolism in *T. gondii* or because of host mitochondria fragmentation [27]. The decrease in taurine was initially intriguing because felines are the definitive host for *T. gondii* and felines are the only mammals that are taurine auxotrophs. While mammals cannot metabolize taurine, several bacterial species can [28-29]. Our initial hypothesis was that *T. gondii* metabolized taurine, but experiments with ^15^N taurine labeling during infection showed that taurine was not metabolized when the ^15^N label was only found in taurine. Infection using metabolomic media, which does not contain taurine, and metabolomics media supplemented with physiological levels of taurine demonstrated that *T. gondii* infection causes taurine secretion in taurine-free media but supplementation lead to no increase in media taurine (Fig S5). Thus, the changes in taurine levels are likely caused by the absence of taurine in the metabolomics media, and not due to parasite metabolism. It is probable that the taurine release is associated with *T. gondii* infection increasing phospholipase A_2_ activity [30-31] as PLA_2_ activity causes taurine secretion [32].

Abundance of multiple amino acids increased during infection, including two amino acids *T. gondii* is auxotrophic for, tryptophan and arginine. Humans are unable to synthesize tryptophan and must import it via LAT1. Transcription of LAT1 increased during infection although it is unclear whether this gene is upregulated directly by *T. gondii* or if *T. gondii* uptake of tryptophan causes the host cell to sense a reduction in intracellular tryptophan levels and thus upregulated LAT1. The increase in argininosuccinate abundance is paired with increased transcription of the rate limiting step in arginine synthesis, argininosuccinate synthetase (ASS1). The cause of increased expression is unknown but ASS1 activity is critical to *T. gondii* replication as previous studies have shown host arginine synthesis is essential to lytic parasite growth [2].

Nucleotide intermediates were broadly more abundant in the infection metabolome, including nucleotides, nucleosides and the essential building block PRPP. *T. gondii* can synthesize pyrimidines but is a purine auxotroph. Host nucleotide metabolism was transcriptionally activated during infection, and key enzymes in *T. gondii* salvage pathways including adenosine kinase (AK) and hypoxanthine phosphoribosyltransferase (HXGPRT) were highly expressed. *T. gondii* putative nucleotide synthesis genes, including phosphoribosyl pyrophosphate synthase (PRS1), were also highly transcribed during infection. Our results support previous findings that *T. gondii* is reliant on host purine synthesis and subsequent salvage, but also confirm that parasite nucleotide metabolism is highly active during infection.

Prior studies have suggested that *T. gondii* possesses a highly active glycolytic pathway and that the parasite may activate host glycolysis through HIF1α activation [33- 34]. Multiple glycolytic intermediates were more abundant in the infection metabolome, including lactate and 2,3-BPG. Activation of glycolysis is driven by both host and parasite through increased expression of multiple key enzymes including HK1, PK1, and LDH in both organisms. The source of increased 2,3-BPG abundance is unknown due to the lack of an identified *T. gondii* BPGM, but the phenotype is intriguing because of the ATP deficit that 2,3-BPG synthesis causes. 2,3-BPG is a known allosteric regulator so it may play a secondary role during infection unrelated to glycolysis [22,35].

Recent publications have greatly expanded our understanding of the *T. gondii* TCA cycle beyond its role in energy production [6,36]. These findings include that the mitochondrial source of acetyl-CoA is from BCKDH [36] and that a GABA shunt allows *T. gondii* to use glutamine to fuel its TCA cycle [6]. Also, *T. gondii* isocitrate dehydrogenase, which catalyzes the rate limiting step of the TCA cycle, is not NAD+ dependent as the human enzyme but instead NADP+ dependent [21]. Multiple TCA intermediates were more abundant particularly late in the infection metabolome, and the paired transcriptome showed that this increase was likely driven by *T. gondii* and not the host. These findings support prior work showing the importance of the *T. gondii* TCA cycle to parasite replication and the inhibition of host TCA activity [6,27].

In contrast to the *T. gondii* TCA cycle, little research has investigated the PPP. The PPP has oxidative and non-oxidative halves and produces ribose-5-phosphate and NADPH. The energetically driven oxidative pathway synthesizes NADPH and funnels carbon from glucose-6-phosphate into the pentose monophosphate pool. From that pool, carbon can either flow into biosynthetic pathways or the non-oxidative pathway. The non-oxidative PPP is catalyzed by transketolase and transaldolase, and is a bidirectional connection between the pentose and hexose monophosphate pools. Previous work has demonstrated that *T. gondii* has an active oxidative PPP [10] and has annotated genes for the transketolase and transaldolase enzymes of the non-oxidative PPP [37]. Metabolomics studies using purified parasites have repeatedly observed PPP intermediates and demonstrated that there is active flux of carbon through the pathway [6,8]. Intermediates in the oxidative and non-oxidative pathways were more abundant in the infection metabolome. The infection transcriptome shows this increased activity is driven by the host and parasite. Host oxidative PPP genes are highly upregulated during infection, while non-oxidative genes are moderately regulated but consistently expressed. The lack of identified oxidative PPP genes in *T. gondii* complicates analysis but there was high expression of putative oxidative PPP genes and non-oxidative genes throughout infection. The infection PPP clearly warrants further study, given the uncharacterized *T. gondii* oxidative pathway enzymes and the unknown cause of increased host PPP activity.

Increases in S7P and SBP abundance, mirrored by increases in O8P and OBP, led us to investigate the link between these metabolites. SBPase was the most likely candidate as it catalyzes the production of S7P and O7P from SBP and OBP, respectively. SBPase activity pulls carbon from glycolysis into S7P in the non-oxidative PPP, where it then can be converted into ribose-5-phosphate. SBPase activity creates an energetically driven non-oxidative PPP by the dephosphorylation of SBP, which may be necessary as the *T. gondii* TCA cycle is NADP+ dependent. While animals do not have SBPase activity, *T. gondii* has a putative SBPase (TGME49_235700). Using a previously published glucose-6-^13^C labeling method, we assayed for SBPase activity. Cells lacking SBPase will generate unlabeled or +1 ^13^C S7P while SBPase produces a pool of +2 S7P. Uninfected cell S7P was either +1 or unlabeled as expected while infected cells had a large pool of +2 S7P, confirming *T. gondii* possesses an active SBPase. SBP and S7P labeling can also be used to estimate SBPase activity, showing that over 90% of S7P in the infection metabolome came from SBPase activity. To confirm SBPase activity, we used a lentiviral system to express TGME49_235700 in HeLa cells along with a BFP negative control and measured SBPase activity with the labeling assay. BFP cells had no significant +2 S7P while cells expressing putative SBPase had a significant +2 S7P pool after only 3 hours of labeling. The complete lack of SBP in the SBPase overexpressing cells, in contrast to the sizable SBP pool in BFP expressing cells, further supports the identification of TGME49_235700 as a SBPase.

To determine the role SBPase plays in *T. gondii* we attempted to delete the gene using SBPase targeted CRISPR/Cas-9 and drug resistance selection inserts for homologous recombination. Multiple attempts yielded no viable knockout parasites. We hypothesized that loss of SBPase reduced the fitness of *T. gondii*. We then tried a rapid selection of entire *Toxoplasma* populations after electroporation using Fluorescence Activated Cell Sorting (FACS) using a positive mCherry and negative GFP fluorescence selections. Untargeted CRISPR/Cas-9 plasmid was used as a negative control. Three independent transformations were sorted; SBPase targeted CRISPR/Cas-9, untargeted CRISPR/Cas-9 and an equal mix of SBPase and untargeted CRISPR/Cas-9. The SBPase targeted transformation generated viable mCherry expressing putative knockout parasites, the untargeted transformation had no mCherry parasites, and the mixed population had half as many mCherry parasites as the SBPase targeted (Fig S6). Putative knockout parasites were cloned, but nine days post-infection, only a few viable plaques grew while most wells contained spacious vacuoles with non-viable parasites. Viable parasites were confirmed as WT SPBase by Southern blot. Although no SBPase knockouts were isolated, these results strongly suggest that viable knockout parasites were present shortly after electroporation, but were unable to grow after FACS sorting and passage. It is likely that deletion of SBPase forces *T. gondii* to produce ribose-5-phosphate exclusively through the oxidative PPP, which creates an imbalance in the NADPH:NADP+ ratio resulting in oxidative stress and parasite death after several rounds of replication. This imbalance would be compounded by the highly active *T. gondii* TCA cycle which also produces NADPH.

To determine the role of SBPase in *T. gondii* metabolism the gene was overexpressed from the α-tubulin promoter. SBPase overexpression led to increased abundance in S7P, several nucleotide metabolites and glycolytic intermediates. This confirms that *T. gondii* SBPase drives carbon into S7P where it can flow into either ribose-5-phosphate and nucleotide metabolism or down the non-oxidative PPP and back into glycolysis. These changes were more pronounced in SBPOE2 than SBPOE1, which was supported by qPCR data showing higher levels of SBPase expression in SBPOE2 than SBPOE1. Both clones were assayed for increased SBPase flux but no changes were apparent. Given that SBPase is responsible for over 90% of S7P synthesis in WT parasites, it is likely that overexpression of SBPase cannot increase SBPase flux, even if the total amount of S7P does increase. SBPase overexpression did result in a significant increase in non-oxidative PPP flux relative to oxidative flux during SBPOE1 and SBPOE2 infection. Together these results demonstrate that SBPase catalyzes a highly active novel metabolic pathway for ribose synthesis via the non-oxidative PPP in *T. gondii*. SBPase likely plays an important role in regulating *T. gondii* ribose synthesis, acting as a switch to funnel carbon from glycolysis into ribose synthesis while avoiding potential redox imbalances that high flux through the oxidative PPP would cause.

## METHODS

### *T. gondii* strains and cell culture

Low passage type II ME49 *T. gondii* used in all experiments. Human Foreskin Fibroblasts (HFFs) from the ATCC were grown in 60mm dishes in DMEM with 10% Fetal Bovine Serum (FBS), 2 mM L-glutamine, and 1% penicillin-streptomycin (Sigma-Aldrich). Once HFFs were in deep quiescence, defined as 10 days post confluency, DMEM media was changed to metabolomic media, RPMI1640 supplemented with 2 mM L-glutamine, 1% FBS dialyzed against PBS (MW cutoff of 10 kD), 10mM HEPEs, and 1% penicillin-streptomycin. After 35 hours, the media was again changed with metabolic media, 1 hour before infection with *T. gondii*.

### Infection Time Course Metabolomics

HFF dishes in triplicate were infected with 2 × 10^6^ tachyzoites, or mock infected with an equal addition of media by volume. At time points 1.5, 3, 6, 9, 12, 24, 36, and 48 hours post infection (HPI), dishes were washed 3x with ice cold PBS, then quenched with 80:20 HPLC grade Methanol:Water (Sigma-Aldrich). Dishes were incubated on dry ice in a −80°C for 15 minutes. Plates were scraped, the solution removed and spun at 2500 x g for 5 minutes at 4°C. The supernatant was removed and stored on ice, then the pellet was washed again in quenching solution and re-spun. Supernatants were combined, dried down under a N_2_ gas manifold, and stored at −80°C.

Samples were resuspended in 150 μL HPLC grade water (Fisher Optima) for analysis on a Thermo-Fisher Vanquish Horizon UHPLC joined by electrospray ionization (negative mode) to a hybrid quadrupole-Orbitrap high resolution mass spectrometer (Q Exactive Orbitrap; Thermo Scientific). Chromatography was performed using a 100 mm X 2.1 mm X 1.7 μm BEH C18 column (Acquity) at 30°C. 15 μL of sample was injected via an autosampler at 4°C and flow rate was 200 μL/minute. Solvent A was 97:3 water/methanol with 9 mM Acetate and 10 mM tributylamine (TBA) with a pH of 8.2 (Sigma-Aldrich). Solvent B was 100% methanol with no TBA (Sigma-Aldrich). Products were eluted in 95% A/5% B for 2.5 minutes, then a gradient of 95% A/5% B to 5% A/95% B over 14.5 minutes, then held for an additional 2.5 minutes at 5%A/95%B. Finally the gradient was returned to 95% A/5% B over 0.5 minutes, and held for 5 minutes to re-equilibrate the column. Peaks were matched to known standards for identification. Data analysis was performed using the Metabolomics Analysis and Visualization Engine (MAVEN) software [38]. Heat maps were generated using the Multi Experiment Viewer program.

### RNAseq

Samples from uninfected or infected 60 mm dishes of HFFs paired to infected metabolomics dishes were resuspended in TRIzol. RNA was extracted per the manufacturer’s protocol, assayed for quality with an Agilent Bioanalyzer and Nano-6000 RNA chip, and synthesized into libraries with a TruSeq RNA Library Preparation Kit v2. Sequencing was performed through the UW Biotechnology Sequencing Center, using Illumina HiSeq2000. Unfortunately the 3 HPI sample was compromised during library preparation and therefore there is no expression data for that time point. The accession number for this data is PRJNA497277 through the NCBI Sequence Read Archive. All *T. gondii* gene accession numbers can be found on ToxoDB.org.

### Read Mapping and Analysis

Approximately 370 million 150 base pair reads were generated from the 9 samples and uploaded to the online Galaxy platform. Reads were filtered using the FASTQ Groomer (version 1.0.4) and then aligned to the *T. gondii* and human genomes (TgMe49 version 2013-04-23, Human version 2013-12-24 HG38) using Tophat2 (version 2.0.14), with the parameters as follows: Max edit distance 2, read mismatch 2, anchor length 8, minimum intron length 70, maximum intron length 500000, max insertion and deletion length 3, number of mismatches allowed 2, and minimum length of read segments 25. Read filtering and mapping are summarized in Table 1. Alignments were converted into FPKM expression values using the Cufflinks tool (version 2.2.1.0).

### Taurine Metabolism Assay

HFFs were grown to deep quiescence in 60 mm dishes, then the media was changed to metabolic media for 35 hours. One hour before infection, the media was change to metabolomic media supplemented with 2 mM ^15^N Taurine (Sigma-Aldrich). Dishes in triplicate were infected with 2 × 10^6^ tachyzoites and metabolites were extracted at 3, 6, 9, 12, and 24 HPI as previously described. Metabolites were extracted from a single time point of triplicate mock-infected dishes at 1.5 HPI to measure taurine uptake from the media. Maven was used to identify metabolite incorporation of ^15^N from taurine metabolism.

### Media Taurine Assay

HFFs were grown to deep quiescence in 60 mm dishes, then the media was changed to metabolic media for 35 hours. One hour before infection, the media was change to one of two metabolomic medias: no taurine or 44μM taurine (Sigma-Aldrich). Dishes were infected with 2 × 10^6^ tachyzoites or mock-infected in triplicate for each media condition. Media samples were taken and metabolites were extracted at 0, 1.5, 3, 6, 9, 12, 24, 36, and 48 HPI. Extraction was performed by adding 80:20 HPLC grade Methanol:Water quenching solution directly to media, incubating samples in a −80°C for 15 minutes and then proceeding with extraction at the first centrifugation step. Maven was used to measure media taurine abundance in each condition over the course of infection.

### Sedoheptulose Bisphosphatase Activity Assay

HFFs were grown to deep quiescence in 60 mm dishes as before, then the media was changed to metabolic media for 35 hours. One hour before infection, the media was change to metabolomic media except using glucose free RPMI1640 supplemented with D-Glucose-6-^13^C at normal glucose concentration (2 g/L) (Sigma-Aldrich). Dishes in triplicate were infected with 2 × 10^6^ tachyzoites, or mock infected with an equal addition of media by volume. At 24 HPI infected and uninfected dishes were quenched and metabolites extracted, processed, and analyzed as previously described, except that samples were resuspended in 75 μL of HPLC grade water and 20 μL was injected. Maven was used to identify the isotopic forms of sedoheptulose-7-phosphate and sedoheptulose-1,7-bisphosphate.

### Sedoheptulose Bisphosphatase Overexpression (SBPOE) in *T. gondii*

The transcript of the putative *T. gondii* sedoheptulose bisphosphatase (TGME49_235700) was amplified from ME49 cDNA made with SuperScript^®^ III First-Strand Synthesis System for RT-PCR (Invitrogen). The product was joined with a *T. gondii* α-tubulin promoter via Splicing by Overlap Extension (SOE) PCR, then ligated into the dihyrofolate reductase plasmid [39] using Gibson Assembly. Low pass ME49 was electroporated with 20 μg linearized plasmid and selected with 1 μM pyrimethamine. Parasites were cloned by limiting dilution and examined by Southern Blot.

### Sedoheptulose Bisphosphatase Overexpression in HeLa cells

Open reading frame (ORF) of the putative *T. gondii* sedoheptulose bisphosphatase (TGME49_235700) was amplified from ME49 cDNA, cloned into pENTR Tev-D-TOPO, then recombined into pLX304 using Gateway cloning. The expression plasmid was transfected in 293T cells with psPAX2 and pCMV-VSV-G and lentiviruses were harvested from the supernatant 48 HPI. 75% confluent 6mm dishes of HeLa cells acquired from the ATCC were infected with 200 μL of lentivirus mixed with 0.2 μL of 8mg/mL Polybrene. 24 HPI media was replaced with media containing 10 μg/mL blasticidin for 3 passages post clearance of transduced cells [40]. As a negative control BFP was inserted into the genome of a second population of HeLa cells using the same delivery system. Both populations were screened for SBPase activity in triplicate at 70% confluency using previously described isotope labeling methods, but without *T. gondii* infection and with three hours of growth in labeled media.

### Pentose Phosphate Pathway Isotopic Flux Analysis

Labeling was performed using metabolomic media made with glucose free RPMI1640 supplemented with D-Glucose-1,2-^13^C_2_ at normal glucose concentration (2 g/L) (Sigma-Aldrich). HFFs were grown to deep quiescence in 60 mm dishes, before being infected with 2 × 10^6^ SBPOE tachyzoites or parental WT ME49 in quadruplicate. At 24 HPI, dishes were quenched, their metabolites extracted and analyzed as previously described. Samples were resuspended in 75 μL of HPLC grade water and 20 μL of sample was injected, to improve peak resolution. Maven was used to identify the isotopic forms of ribose-5-P.

### qPCR

Primers specific to *T. gondii* Tubulin subunit A or human actin subunit B were used as the control. SBPase expression was compared between *T. gondii* strains SBPOE1, SBPOE2 and WT, and between HeLa populations with SBPOE or with BFP. cDNA was extracted from all cell types using TRIzol and converted to cDNA as previously described. qPCR was performed using iTaq Universal SYBR Green Supermix.

## Acknowledgements

We would like to thank Anthony Dawson for generating the lenti viruses and assistance with expression protocols, Jing Fan for her advice on project set up and Amy Caudy for her advice on carbon isotope labeling and the sedoheptulose bisphosphatase assay.

**S1. Representative spectra demonstrate SBPase activity in infected samples** Representative spectra of S7P from uninfected (top) and infected (bottom) cells grown with glucose-6-^13^C (M= parental unlabeled, M+1 = mono C13 labeled, M+2 = double C13 labeled, M+3 = triple C13 labeled). M+2 label demonstrates the presence of SPBase activity.

**S2. SBPase and BFP expressing HeLa fluorescence:** Differential Contrast and fluorescent images in the blue channel from SBPase (A and C) and BFP (B and D) expressing HeLa cells respectively. Lentiviruses were used to stably incorporate either SBPase or BFP into HeLa genomes. Fluorescence microscopy confirmed BFP expression in the BFP population but not in the SBPase cells. qPCR confirmed SBPase expression was only present in the SBPase expressing cells and not the BFP cells. Scale bars are 100 μm.

**S3. SBPOE1 and SBPOE2 southern blot:** Southern blotting identified two unique SBPOE clones. Genomic DNA was extracted from clonal populations of *T. gondii*, digested with restriction enzyme and assayed via Southern blot with an SBPase targeted probe. A wild type band was present in all populations and a secondary band was observed in SBPOE1 and SBPOE2, indicating a second SBPase gene insertion. The secondary band sizes were different in SBPOE1 and SBPOE2, proving they are unique clones. Gel displays ladder (L), SBPOE1 (1), SBPOE (2), and wild type (W) from left to right.

**S4. Ribose-5-phosphate was differentially C13 labeled during SBPOE1, SBPOE2, and WT infection**

SBPOE1, SBPOE2, and WT infected cells were grown in quadruplicate with glucose-1,2-^13^C for 24 hours. Ribose-5-phosphate isotope composition was determined using HPLC-MS and Maven as previously described.

**S5. Taurine media abundance:** Triplicate infected and uninfected media samples were taken over the 48 hour time course from dishes in standard metabolomic media (0 Taurine) or from media supplemented with taurine (44 μM Taurine). Metabolites were extracted and quantified with HPLC MS. Infection media taurine abundance was averaged and normalized to the average uninfected media abundance then log base 2 transformed. In standard metabolomic media with no taurine infection causes a rapid secretion of taurine which levels off at 24 HPI. Media supplemented with taurine at roughly physiological levels shows no taurine secretion.

**S6. FACS analysis of attempted SBPase knockout with mCherry positive selection:** Fluorescent particle analysis of sorted *T. gondii* populations. Three *T. gondii* populations were electroporated with an mCherry positive selectable marker and one of three CRISPR/cas-9 plasmids; SBPase targeted CRISPR/Cas-9, untargeted CRISPR/Cas-9, and an equal mix of SBPase and untargeted CRISPR/Cas-9. Electroporated parasites recovered and grew for 72 hours before FACS sorting and cloning. The SBPase targeted CRISPR population yielded viable mCherry expressing parasites, the mixed CRISPR population had half as many viable mCherry positive parasites, and the untargeted CRISPR population had no mCherry expressing parasites.

